# Cross-modal Semantic Relationships Guide Spontaneous Orienting in Real-life Scenes

**DOI:** 10.1101/2021.11.29.470351

**Authors:** Daria Kvasova, Travis Stewart, Salvador Soto-Faraco

## Abstract

In real-world scenes, the different objects and events available to our senses are interconnected within a rich web of semantic associations. These semantic links help parse information and make sense of the environment. For example, during goal-directed attention, characteristic everyday life object sounds help speed up visual search for these objects in natural and dynamic environments. However, it is not known whether semantic correspondences also play a role under spontaneous observation. Here, we investigated this question addressing whether crossmodal semantic congruence can drive spontaneous, overt visual attention in free-viewing conditions. We used eye-tracking whilst participants (N=45) viewed video clips of realistic complex scenes presented alongside sounds of varying semantic congruency with objects within the videos. We found that characteristic sounds increased the probability of looking, the number of fixations, and the total dwell time on the semantically corresponding visual objects, in comparison to when the same scenes were presented with semantically neutral sounds or just with background noise only. Our results suggest that crossmodal semantic congruence has an impact on spontaneous gaze and eye movements, and therefore on how attention samples information in a free viewing paradigm. Our findings extend beyond known effects of object-based crossmodal interactions with simple stimuli and shed new light upon how audio-visual semantically congruent relationships play out in everyday life scenarios.

## Introduction

Real world environments are characterized by their high complexity. The efficiency with which humans sample visual information in such complex visual scenes poses a problem to traditional theories of visual search that are heavily based on feature analysis. Recent accounts have proposed that higher level processing, based on attentional templates that incorporate patterns and object-specific categories in the visual cortex, can guide information sampling efficiently in everyday real-world scenes. These attentional templates harness on statistical regularities learned throughout experience, familiarity with certain visual patterns, predictable functional relationships between objects, and contextual constrains (e.g., Papeo et al. 2019; Peelen & Kastner, 2015; Soto-Faraco et al., 2019). However, one important feature of real-life environments is that information arrives from a variety of sensory channels. Therefore, it is important to understand whether, and how multisensory information is used during sampling in complex environments.

Cross-modal interactions have been under the focus of research for various decades. In addition to well-known cross-modal spatial and temporal effects on attention orienting (e.g., Spence & Driver, 1994; 1998; McDonald et al., 2000Van der Burg et al., 2008; Van den Brink et al., 2014; see Spence & Soto-Faraco, 2019 for an applied perspective), other lines of research have started to unveil the potential of cross-modal semantic information. Behaviorally relevant information about the identity and semantic attributes of objects, and their semantic correspondences can also help us parse sensory information and deploy processing resources in an efficient manner (Iordanescu et al., 2008, 2010; Mastroberardino et al., 2015; Kvasova and Soto-Faraco, 2019).

Many of these prior studies, however, have typically used simplified displays which allow for optimal experimental control but may fail to capture the complexity of relevant real-world scenes (e.g., Soto-Faraco et al., 2019; Matusz et al., 2019, for a similar argument). To overcome this problem, a recent study by Kvasova et al. (2019) addressed the generalization of cross-modal semantic facilitation, using a search task in real-life scenes presented on video. Despite this generalization from simple to complex scenes is important, participants sampling was motivated by an explicit search task. Therefore, this finding is unrepresentative of situations of spontaneous viewing, when the observer has no specific goal in mind, or the goal is unrelated to finding a particular target object in her environment. Here, we addressed the potential of cross-modal semantic cues to guide the deployment of spontaneous spatial attention under conditions in dynamic scenarios containing real-life complexity.

Previous studies have demonstrated cross-modal semantic congruence effects on visual processing and attention. For example, Molholm et al. (2004) found better performance for audio-visual object recognition compared to unimodal performance, when the auditory and visual semantic cues were congruent. Similar cross-modal object recognition improvements were seen with haptic-visual stimuli by Pesquita et al. (2013). Cross-modal semantic congruence can also improve visual detection (Chen and Spence, 2011), and picture naming (Mädebach et al., 2017) and, especially relevant for the question at stake here, cross-modal semantic congruence influences the spatial distribution of visual attention (Iordanescu et al., 2008; 2010; Mastroberardino et al., 2015; Kvasova et al., 2019). Despite these findings strongly suggest that semantic information plays a role in spatial orienting, most of these outcomes have been observed using a reductionist approach which trades ecological validity for experimental control (for a similar argument see, Blanton & Jaccard, 2006; Burgess et al., 2006; Kayser, Körding, & König, 2004; Kingstone et al., 2003; Neisser, 1976; 1982; Soto-Faraco et al., 2019). For example, the studies mentioned so far have used simplified scenarios in which isolated visual objects are presented in the absence of any meaningful or structured context.

The role of meaning and context on the guidance of attention have recently come under the spotlight of visual attention research, highlighting important the differences between simple meaningless artificial search displays and naturalistic scenes (Peelen et al., 2014; Henderson and Hayes, 2017). For instance, humans can extract complex information from just a brief glance at a scene, and can then predict which types of objects, which likely spatial arrangements, and what semantic connections between objects are likely to be found (Wu et al., 2014). In addition, real world scenes are typically characterized by high perceptual load, compared to simplified search arrays (Soto-Faraco et al., 2019). Given these differences, it would be fair to assume that visual search and eye movement behavior within a complex scene might be significantly different from that of simple search paradigms used in most previous studies. Hence, generalization studies are warranted.

So far, generalization studies addressing attention orienting in complex, realistic scenarios, have mostly focused on visual-only tasks. Based on the scarce previous literature it is difficult to know if laboratory results addressing cross-modal semantic effects on orienting generalize to complex real-life-like scenes. In one of the few studies to address this question, Nardo and colleagues (2014) measured eye movements and BOLD responses using fMRI whilst participants passively watched video-clips of real-life scenes. Nardo et al. reported that semantic congruence between sounds and visual events did not produce any modulation in gaze distribution or brain activations (despite other aspects of the stimuli, such as spatial congruence between visual and acoustic cues did). This null effect of semantic congruence in realistic scenes contrasts with the significant outcomes from various, more controlled laboratory studies discussed earlier (Iordanescu et al., 2008; 2010; Mastroberardino et al., 2015). One could think that this constitutes a case in which laboratory findings magnify an effect that is not very significant in real life behavior. The reduced impact of semantic congruence whilst freely observing real scenes in Nardo et al.’s study could be due, for example, to the presence of more preponderant spatial and temporal audio-visual cues, which overrode potential cross-modal effects based on object identity.

A recent visual search study from our group (Kvasova et al., 2019), also using realistic video clips, found that semantically consistent but spatially and temporally uninformative sounds speeded up visual search times, in comparison to distractor-consistent, neutral sounds, or no sound conditions. However, unlike Nardo et al., Kvasova et al.’s study used a goal-oriented search paradigm. Given that in Nardo et al.’s experiment, subjects did not have a specific task other than to view the video clips, one could think cross-modal congruence simply is inefficient when task irrelevant. Whilst Mastroberardino and colleagues (2015) demonstrated that audio-visual semantically congruent events can indeed influence and attract attention despite being unrelated to the task, in that study stimuli were not realistic scenes.

The combination of results discussed above, and the substantial methodological differences between studies, leaves the main question unresolved. Indeed, one is left to wonder whether semantic coincidences across modalities in everyday life scenes do exert some attraction effect on spontaneous attention. That is, even if not necessarily relevant to our current goals. Here, we use eye-tracking to investigate the impact of cross-modal congruence on the spontaneous direction of attention as people simply watch real world scenes. In contrast to Nardo et al., we focused on scenes where there are no salient spatio-temporal cues, and therefore semantic cues may become more relevant. Using eye-tracking allowed us to test participants as they freely watched the scenes and orienting unfolds spontaneously. In particular, we tested whether semantically congruent sounds produce faster, longer and/or more frequent overt spatial orienting towards their affiliate visual object in real-life dynamic scenes.

Each participant was presented with short video-clips of everyday life scenes with background noises. We defined an area of interest (AOI) for each video, containing a specific object of interest. Each video was dubbed with three different soundtracks defining three different conditions: *consistent* (a characteristic object sound consistent with the object of interest), *neutral* (a characteristic object sound not corresponding to any object in the scene) and *no characteristic sound* (videos are presented only with the background noises). We hypothesized that semantically consistent sounds would increase the salience of the corresponding visual object, and therefore the probability of directing overt attention toward the visual object would increase. In order to test this hypothesis, we measured various eye-gaze parameters. First, we computed the probability of looking into the AOI in each condition. Based on our hypothesis we predicted higher percentage of videos where AOI is looked at in consistent than in neutral or no sound conditions. Second, we hypothesized that consistent semantic information would strengthen the effect of spatial orienting and increase exploration of the corresponding visual object. Therefore, we predicted higher total time spent looking (dwell time) and number of fixations inside the AOI in the consistent condition in comparison to the neutral or no sound conditions. Further, we measured how fast the gaze goes towards the AOI. We predicted that in the consistent condition gaze would be directed in the AOI faster in comparison to control conditions.

## Methods

### Participants

Forty-five subjects (12 males, mean age 23.3 (between 19-35) participated in the study. All gave written consent to take part in this experiment. The protocol was approved by the ethical board CIEC Parc de Salut Mar, UPF. All participants had normal or corrected to normal vision, normal hearing and were naïve about the purpose of the study. Each participant performed a calibration run with the eye-tracker software before each experiment block to ensure that accuracy of gaze tracking was acceptable (within <0.5º of visual angle). Participants were excluded if calibration failed to reach the criterion after 3 attempts. Sample size was set a priori, but without any estimation of statistical power.

### Stimuli

We created a set of 108 video clips of 2s duration selected from movies, television, YouTube or recordings by the authors. All videos, size 1024×768 pixels and 30 fps were edited with iMovie software 10.1.10. We replaced the original soundtracks with background noises composed of generic sounds typical of the corresponding visual scene. The acoustic background noise was introduced to avoid possible cross-modal spatio-temporal correspondences present in the original soundtracks, and to attenuate the alerting effect of the characteristic object sounds used in the conditions of interest. Background noise was tailored to the videos. For example, if the video contained scenes from a concert, we added a noise of a crowd (see example video clips and sounds in the online supplementary materials).

Each video contained a designated target object (e.g.: a guitar, a bicycle, a dog, etc.). Target objects were chosen so that they were visible throughout the clip (no occlusions, good contrast), remained in a relatively stable position, were not part of the main action, and were not placed at the center of the frame. These criteria were checked thoroughly by at least two judges on each video. We counterbalanced the videos across the conditions of interest and participants (see below), so that each video contributed in all conditions in equal proportions in the final dataset. This should compensate for intrinsic variability of the materials unrelated to the conditions of interest. We defined an area of interest (AOI) in each videoclip, around the object of interest. Due to the high heterogeneity of object’s size and location, the areas of interest were defined manually for each clip by creating a rectangular area that encompassed the object in all frames. This method of AOI definition has proved useful when dealing with complex and heterogenous visual scenes (Hessel et al., 2016, for a review of AOIs methods).

Characteristic object sounds were obtained from Freesound.org database. When present, the object sounds provided no information about the location of the object (as they were always presented from the same, central, location) or its temporal profile (the sound onset was fixed for all videos and conditions). All the sounds were normalized and presented at a comfortable intensity. Object sounds, when present, where +10dB SPL compared to the background noise, and their duration was variable due to differences in their profile (M = 600 ms, SD = 145 ms). Sounds were delivered through two loudspeakers placed at each side of the monitor, in order to render them perceptually central.

The design contained three sound conditions. In the *no sound condition* the video soundtrack contained background noise suitable to the content of the scene but no discernible characteristic object sounds. In the *consistent condition*, the characteristic sound of the object of interest of that video was embedded in the background noise (e.g., the sound of a barking dog when a dog is the object of interest). In the *neutral condition*, a sound characteristic of an object not present in the video was embedded in the background noise (e.g.: the sound of a piano when a dog is the object of interest, and the scene doesn’t contain a piano).

Please note that objects of interest, and therefore the characteristic sounds, could repeated across different videos (different dogs, different guitars, etc.). To prevent spurious semantic effects in the neutral sound condition we used the addition criterion that the sounds in this condition could not belong to the same semantic category as the object of interest (e.g.: a video where a dog is the object of interest could not be presented with a sound of a piano in the neutral condition). We implemented this rule automatically by using five semantic groups (animals, vehicles, electronics, musical instruments and, other), and then we inspected the assignments manually and if some kind of obvious semantic association was found, the sound was reassigned.

The experiment was programmed and conducted using the Psychopy package 1.84.2 (Python 2.7) running under Windows 7. Participants were sitting in front of a 22.5” computer monitor (Sony GDM-FW900) at a distance of 70cm. Two loudspeakers, delivering the sounds stereophonically, were placed at each side of the monitor. Eye movements were recorded using the EyeTribe eyetracker (60 Hz sampling rate and 0.5° RMS spatial resolution) and PyGaze 0.5.1 open-source software.

### Procedure

The video-clips were counterbalanced across conditions and participants by creating three equivalent versions of the experiment, each presented to N=15 participants. Each participant was presented with 108 unique trials, divided into six blocks of 18 videos (six videos per condition per block). The order of blocks and videos within blocks were randomized for each participant. Overall, each video-clip once appeared in each condition the same number of times across the experiment.

Participants were instructed to simply watch the videos as if they are watching television, with the only constrain to watch within the screen area (e.g., avoid looking aside of the screen or closing the eyes). This way we attempted to minimize any task related confound. The eye-tracking equipment and sensors were calibrated individually for each participant and before each of the six blocks of the experiment.

Each trial (see Figure 1A) began with a central fixation cross in a blank screen. Participants had to fixate on the cross for the trial to start by moving the circle that represented their gaze position to the cross. Once gaze was detected in the area of fixation cross the trial began. The trial consisted of the 2s video with the corresponding soundtrack. When a characteristic sound was presented (consistent or neutral conditions), its onset preceded the videoclip by 100ms. After the videoclip, a 1s blank screen preceded the next fixation or end of the block. Participants were encouraged to rest between blocks, but they could take additional self-paced rest periods by not fixating on the cross between trials. We recorded eye position from the onset to the end of each videoclip.

**Figure 1.**
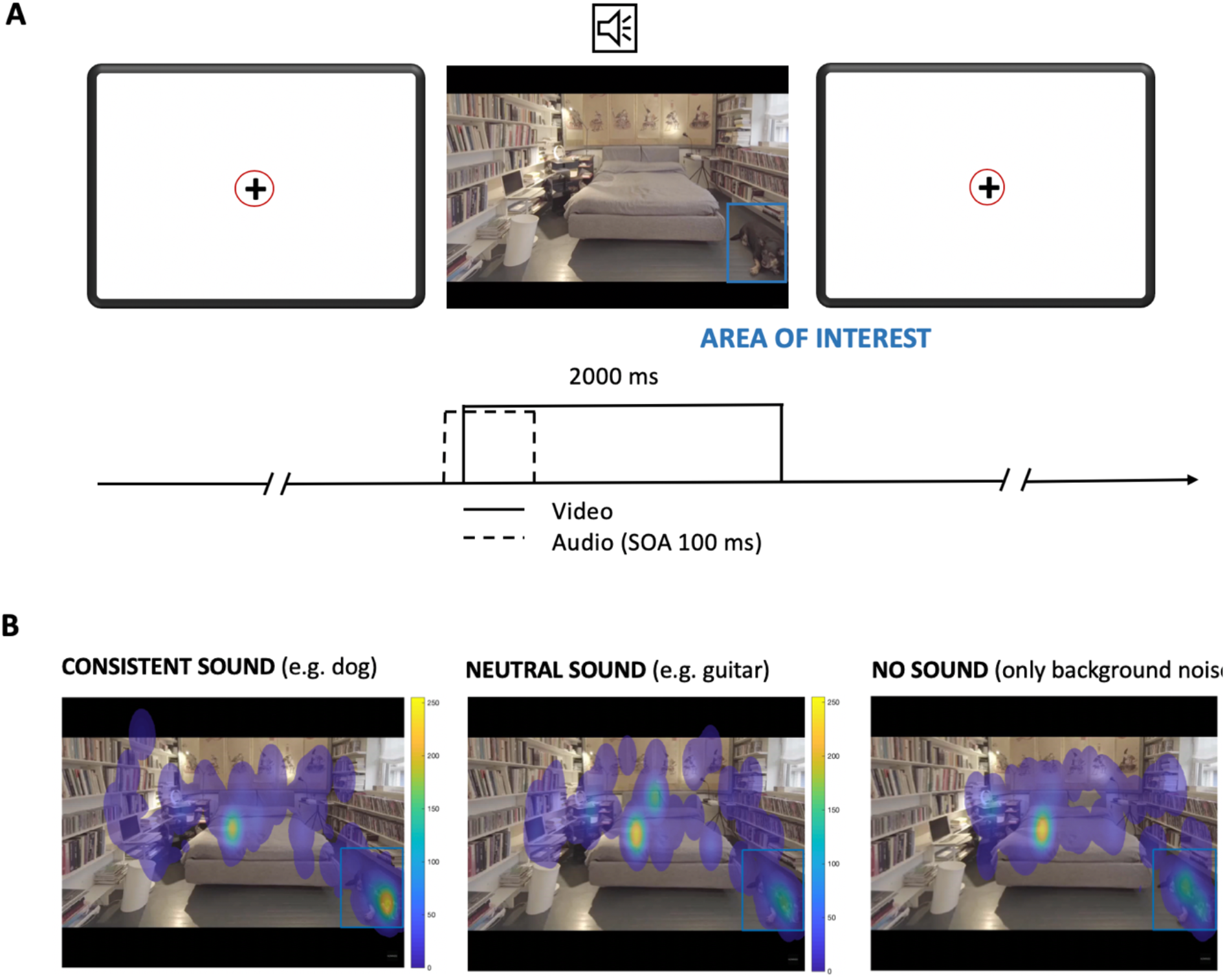
(*A)* Sequence of events in a trial. The trial started with the presentation of a fixation cross. Participants were asked to fixate on the cross, moving the circle (representing their gaze position) to the cross. Once the participant was fixated on the center of the screen the trial started. In the *consisten*t and *neutral* condition, the trial started with the sound that was either characteristic to the one of the objects (*consistent*) or not characteristic to any object on the video (*neutral*). Participants were instructed to fixate on the cross but once the video started, they could move their eyes and freely watch the video. The blue rectangle outline indicates the area of interest containing the object of interest (this marker was not presented to the participant). Here it is placed on top if the video for the illustrative purposes. (*B)* The results of one single example videoclip. The color heat indicates fixation behavior averaged over each condition (15 participants per condition). This visualization displays the distribution of gaze overlaid on one representative frame of the videoclip. Color heat (from yellow to blue) represent in descending order the amount and duration of fixations on each screen area, with ratio from 0 to 250.

## Results

We performed detection of fixations in the Area of Interest (AOI) on the raw eye-gaze data using MATLAB_R2017a. Fixation was defined as a period of stable gaze within 1º of visual angle with duration >50 ms. We calculated four measures: percentage of videos where the AOIs was fixated, number of fixations inside AOI, dwell time inside the AOI, and time to first fixation inside AOI. Dwell time and time to first fixation analyses were conditioned to at least one fixation inside the AOI having occurred. All the measures were calculated for each participant and condition, and then averaged across participants.

For illustrative purposes, we generated heat maps to visualize the general distribution of gaze points in all experimental videos in the three sound conditions (see an illustrative example in Figure 1B). We performed data analyses using a repeated measures ANOVA separately on each of the four measurements listed above, with subject as the random effect and auditory condition as the factor of interest. The results are illustrated in Figure 2.

**Figure 2.**
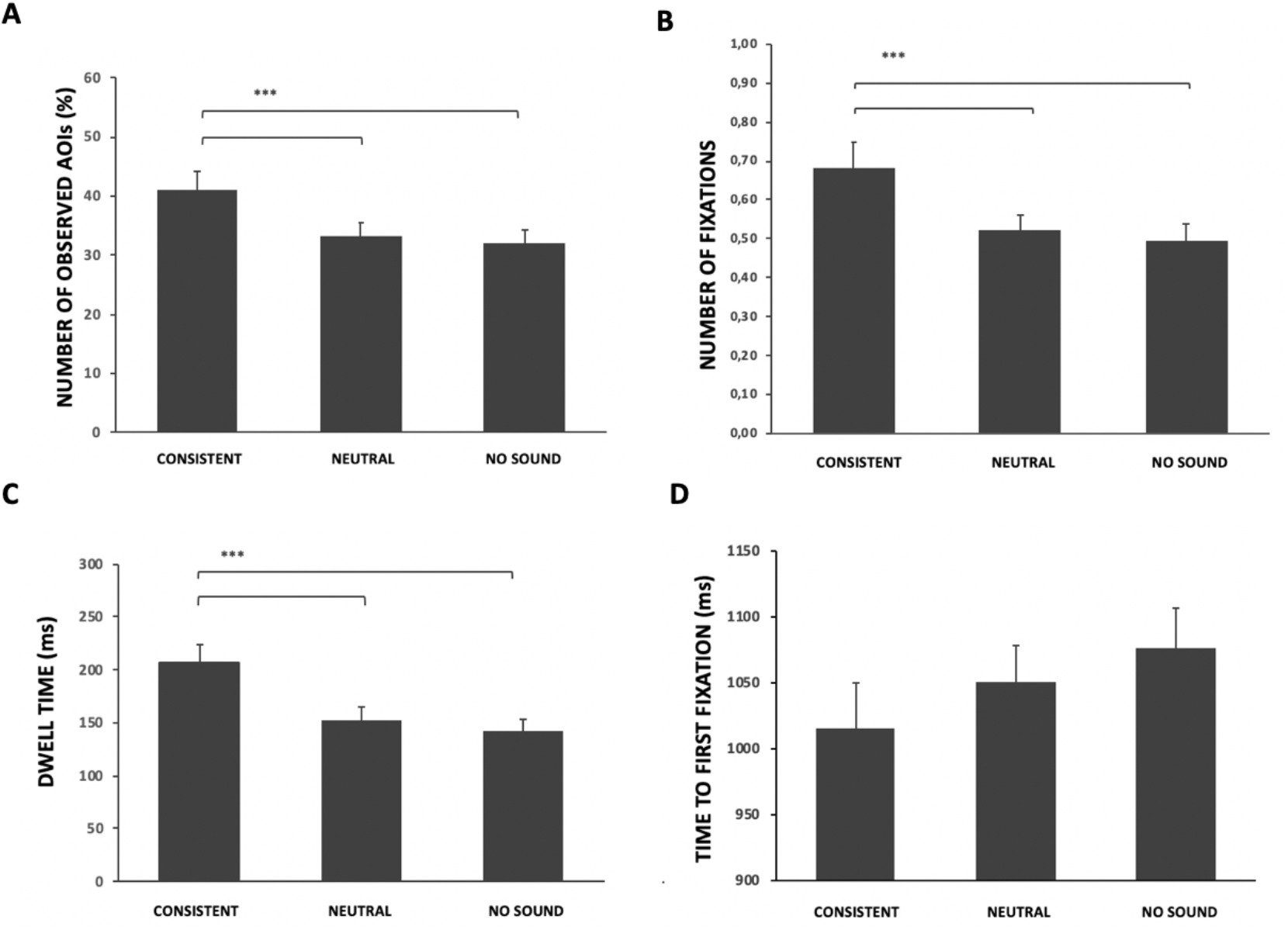
*(A)* The percentage of videos where participants looked at the area of interest. (B) Number of fixations detected inside of the region of interest. (C) Total dwell time spent looking inside of the area of interest. (D) Latency of first fixation inside the area of interest. Error bars indicate the standard error. * p-value<0.05, **p-value<0.01, ***p-value<0.001.

The analysis on the percentage of videos with a fixation in the AOIs returned a significant main effect F(2,88)=14.37; p=0.000004. Based on the significant main effect, we went on to test our specific prediction using t-tests. In keeping with the hypothesis, the analysis demonstrated that the percentage of videos where the AOI had been fixated was higher in the consistent condition M=41% than in the neutral M=33% [t(44)=2.22, p=0.00015, Cohen’s d=0.45] and no sound M=32% [t(44)=2.89, p=0.00002, Cohen’s d=0.5] conditions (*Figure 2A*). In summary, an object was 9% more likely to be looked at if its characteristic sound was played.

The analyses on the number of fixations inside the AOI F(2,88)=10.91; p=0.00006 and dwell time inside AOI F(2,88)=17.23; p=0.0000005 also returned significant effects. (These two variables are logically correlated). We had hypothesized that consistent auditory semantic information will strengthen spatial orienting towards visual object and increase exploration inside the AOI. Therefore, we predicted that dwell time and number of fixations inside the AOI in the sound-consistent condition would be higher in comparison to the neutral or no sound conditions. One-tailed t-tests showed that number of fixations was higher in the consistent condition (M=0.68) in comparison to neutral (M=0.52), [t(44)=3.12, p=0.0016, Cohen’s d=0.44] and no sound (M=0.5), [t(44)=4.22, p=0.00006, Cohen’s d=0.5] conditions (*Figure 2B*). Accordingly, dwell time was longer in the consistent condition (M=207 ms) in comparison to neutral (M=153 ms), [t(44)=3.94, p=0.0016, Cohen’s d=0.55] and no sound (M=143), [t(44)=5.01, p=0.00006, Cohen’s d=0.66] conditions (*Figure 2C*). All the above comparisons are one-tailed (given the directional hypothesis) and survived the multiple comparison correction using Holm-Bonferroni (Ludbrook, 1998).

Finally, the analysis of time to first fixation did not return significant results F(2,88)=1.31; p=0.27. Because our last hypothesis stated that semantically consistent sound will guide attention to the visual object faster than neutral or no sound conditions, we had predicted that participants would look into the AOI sooner in the consistent condition than on the other conditions. This did not happen. Further exploratory t-tests showed only marginal differences in the expected direction (M=61ms) between consistent sounds and no sound [t(44)=1.49, p=0.072, Cohen’s d=0.16] conditions (*Figure 2D*).

## Discussion

This study addressed whether semantic correspondences between visual objects and their sounds influence spontaneous orienting during free viewing of real-life dynamic scenes. Our results suggest that cross-modal semantic congruence indeed has an influence on gaze behavior and on the distribution of spatial attention, in free viewing conditions. Visual objects in control conditions, with neutral sounds or no sound, were looked at only in approximately 32% of times the video was shown. Hearing the characteristic object sound increased the probability that the affiliate visual object will be looked at in the scene to 41%. That is, a random object in the scene is 28% more likely to be looked at, if its characteristic sound is being played. Please note that we intentionally chose videos where the objects of interest had relatively low salience in the image, hence they did not always attract gaze to start with. We also observed that cross-modal semantic congruence increased the number of fixations on the object by around 30% and total dwell time spent looking at the object of interest (inside area of interest) by more than 35%. Despite a trend in the expected direction the analyses did not support that there was a significant speedup of fixations to objects paired with semantically congruent sounds. Overall, however, these results clearly indicate that characteristic sounds might increase the salience of corresponding visual object and drive spatial orienting towards them, even in free viewing conditions of naturalistic complex scenes.

Many studies have investigated the potential benefits of cross-modal congruence using a variety of low-level attributes such as spatial and temporal cues (e.g. Bolognini et al., 2005; Koelewijn et al., 2010; McDonald et al., 2000, 2001; Vroomen & de Gelder, 2000), and higher-level attributes such as semantic correspondences (Chen and Spence, 2011; Iordanescu et al., 2008, 2010; Molholm et al., 2004; Pesquita et al., 2013). As mentioned in the introduction, previous findings have shown a facilitation effect of cross-modal object-based congruence on visual search (Iordanescu et al., 2008;2010; Knoeferle et al., 2016; Kvasova et al., 2019). However, in these studies, participants were given the goal to find objects and therefore, actively used the semantic cues for the task. Here, we have shown that these cross-modal semantic effects do have an impact on spontaneous viewing behavior under no, or very weak, task constrains.

The present result has implications about the degree to which voluntary attention guides cross-modal semantic effects on orienting. For example, in the study of Iordanescu (2008) it was demonstrated that semantically congruent sound attracts attention towards predefined visual targets, but not to distractor objects. The authors then claimed that cross-modal facilitation can occur only in goal-directed manner, meaning that sounds can only enhance visual representations if an attentional template is activated for a visual target search. This would imply a strong role of voluntary attention. This finding was further supported by our previous study (Kvasova et al., 2019) where we demonstrated that cross-modal semantic effects extrapolate to search tasks into real-life scenes (as opposed to ordered search arrays). Here, the result of Nardo et al. (2014) is also relevant, given that in their experiment using eye-tracking and free observation found no particular effects of cross-modal semantic congruence on orienting. This would support the same idea, namely, that cross-modal semantics matters only when for goal relevant objects, but wanes when they are task irrelevant. Yet, the present results, under a different set of constrains, point to a modified conclusion. Namely, the present study showed conclusively under free-viewing that observers do orient more frequently and for longer time to objects whose characteristic sound is playing, even if they are not searching or have been primed in any other way. The potential for cross-modal congruence to attract attention even when task irrelevant has been also pointed out by other studies (Mastroberardino et al., 2015; Kvasova and Soto-Faraco, 2019). In both these studies audio-visual objects appeared as task-irrelevant objects before or in parallel with a different primary task. In the two studies, the main finding was that under certain conditions the position of the irrelevant cross-modal congruent objects attracted spatial attention and influenced performance in the primary task.

Hence, previous results seemly point to opposite conclusions regarding the question of whether cross-modal semantic congruence has an effect on spontaneous spatial orienting in real life. How can we reconcile these findings? In the light of the present evidence, it would seem that cross-modal semantic congruence effects on orienting do not abide to a strictly automatic or mandatory process, yet under some conditions semantic links can percolate behavior even if task irrelevant. We believe that the relevant focus of the question is not whether or, but under which conditions cross-modal semantic congruence is likely to impact orienting. For example, previous studies have varied widely in terms of perceptual load, which seems to be a determining factor for whether irrelevant information is influential on behavior or not (Lavie, 2005). This trade-off between perceptual load and processing of task-irrelevant information has recently been applied to cross-modal contexts, using simplified displays. For example, Lunn et al. (2019) showed that cross-modal attentional capture, using simple stimuli, is sensitive to perceptual load (Lunn et al., 2019), and Kvasova and Soto-Faraco (2019) have shown that this perceptual load modulation also applies to cross-modal semantic congruence effects in visual search.

However, perceptual load is not the only relevant variable to explain whether cross-modal semantic effects will occur or not when freely viewing natural scenes. In fact, the high perceptual load typical of the crowded, dynamic natural scenes did not eliminate the impact of cross-modal congruence in the present study. In natural scenes, the impact of the complexity of visual displays may not increase perceptual load in the same way as in artificial arrays of visual objects (Peelen & Kastner, 2011). This is because real-world scenes are meaningful, contain context and learned ‘typical’ structural relationships between elements. For example, cats are seen resting on top of pillows more often than the other way around. These constrains are not usually respected in artificial arrays (Kaiser et al., 2014). It was demonstrated previously that not only low-level visual salience but also semantic relationships between the objects in a complex visual scene can guide attention effectively (Wu et al., 2014, for review). In particular, Henderson and Hayes (2017) showed that while calculating salience of visual scenes one must take into account not only low-level features but high-level object and context information, since both low-and high-level information participate in guiding attention.

These structural properties of natural scenes apply to the multisensory case investigated here, and could underlie the outcome that, despite the perceptual load, cross-modal congruence still played a role in task-irrelevant conditions. This constitutes an example of how the outcomes of laboratory experiments with simplified set ups might vary as conditions approach real life (Soto-Faraco et al., 2019; Matusz et al., 2019; Maguire et al., 2012).

In natural scenes, cues regarding low level features (such as the location of a sound and a visual event) as well as the higher-level features regarding structural correlations or semantic relationships are all present at the same time. Nardo et al. (2014) already demonstrated that low-level spatial correspondences between sounds and visual objects affect spontaneous spatial orienting in real-life scenes. Yet, they did not find any effect of semantic congruence. We reason that the low-level properties in that study overwhelmed the possibly weaker effects of semantic congruence. In our study we minimized the role of low-level cross-modal congruence by making spatial and temporal properties equivalent across conditions. Admittedly, this is unrepresentative of natural audio-visual events, but a necessary measure in order to isolate the semantic effects from any direct attention-grabbing effect of spatio-temporal correspondences.

In terms of the underlying mechanism producing these effects, we have assumed that increased saliency of visual objects by congruent sounds would predict the effects found, as well as a speed up of the latency of first fixations on the object. Given the trend in the expected direction of the latency test, this saliency hypothesis is supported. However, one potential alternative explanation is that the sound of a characteristic object will increase the variability of spontaneous looking. That is, when we hear a recognizable sound, we will tend to examine more. This increased exploration hypothesis could explain all the effects found, given that by exploring more one would have more chances to bump into the object corresponding to the sound, and then stay on it or return to it more frequently thereafter. Indeed, such an account does not predict faster latencies on first fixation in the sound-consistent condition compared to the sound-neutral condition, as both these conditions should be totally equivalent until the object is finally found. Given the lack of difference between the latencies of first fixation between these two conditions, it is not possible to attribute between the increased saliency and the increased exploration accounts, at present.

## Conclusions

In the current study, we examined whether crossmodal sematic congruence guides attention when viewing real-life scenes. We found that characteristic sounds increase the probability to look at the corresponding visual objects and increase total time spent looking and number of fixations at the object of interest. All in all, the results demonstrate that cross-modal semantic congruency can play a role when watching everyday life scenes.

